# Antioxidant properties of *Rhodiola rosea*

**DOI:** 10.64898/2026.05.21.726678

**Authors:** Drew F. Brink, Tiara L. Sapp, Trwska S. Ghafoor, Paige A. Boyland, Yukako C. Tamazawa, Guneet Kaur, Nataliia V. Shults, Robert D. Sullivan, Yuichiro J. Suzuki

## Abstract

*Rhodiola rosea* is a traditional medicinal plant often classified as an adaptogen, with reported effects in supporting the body’s response to physical, environmental, and emotional stressors. The present study investigated the antioxidant properties of *Rhodiola rosea* extract and its major chemical constituents to provide insight into their potential mechanisms of action. Through *in vitro* biochemical assays, we demonstrated that *Rhodiola rosea* extract has the capacity to reduce hydrogen peroxide (H_2_O_2_) levels. Among its primary chemical components, rosavin significantly decreased H_2_O_2_, whereas salidroside had no effect. Neither compound affected superoxide levels. Structural analysis revealed that the intact phenylpropanoid glycoside architecture of rosavin is required for activity, as its individual components, arabinose and rosin, showed no inhibitory effect. Further investigation demonstrated that rosavin attenuates H_2_O_2_-mediated oxidation of thiol groups, supporting a role in cellular redox regulation. In cultured human cells, rosavin mitigated reductions in cell viability induced by exposure to H_2_O_2_, indicating cytoprotective effects under oxidative stress conditions. Finally, in an *in vivo* model, administration of SARS-CoV-2 spike protein increased circulating levels of H_2_O_2_, which were subsequently reduced following rosavin treatment. Collectively, these findings identify rosavin as a structurally dependent antioxidant component of *Rhodiola rosea* that modulates H_2_O_2_-associated oxidative stress and supports further investigation of phenylpropanoid glycosides as adaptogens.

## Introduction

*Rhodiola rosea*, commonly known as “roseroot” or “golden root,” is a perennial flowering plant in the Crassulaceae family whose medicinal properties are derived from its root (Chiang et al., 2015). *Rhodiola* species are commonly classified as adaptogens and have been historically used to enhance resilience to physical and psychological stress and to treat fatigue, infection-related weakness, and chronic illness [Chiang et al., 2015]. Reported pharmacological effects include anti-inflammatory, antioxidant, antidepressant, antimicrobial, immunomodulatory, and cardio-, neuro-, and hepatoprotective activities, along with central nervous system stimulation and enhanced work capacity [Chiang et al., 2015]. *Rhodiola rosea* grows in high-altitude, subarctic environments characterized by low temperatures and harsh environmental conditions, including mountainous regions throughout Europe, Asia, North America, Britain, and China [Liang et al., 2024]. The medicinal properties of *Rhodiola rosea* are primarily attributed to its diverse group of bioactive constituents, including phenolic glycosides, polyphenols, organic acids, and essential oils [Chiang et al., 2015]. The principal compounds associated with its adaptogenic and pharmacological effects are the phenylethanoids and phenylpropanoids, such as salidroside, tyrosol, rosavin, rosarin, rhodionin, rhodiosin, and rosin [Booker et al., 2016; Chiang et al., 2015]. Commercial preparations are typically standardized to 1% salidroside and 3% rosavin; however, the content of these active compounds varies widely due to environmental and processing factors such as geographic origin, climate, harvest conditions, and extraction or drying methods, contributing to batch-to-batch variability and potential differences in therapeutic effects [Dimpfel et al., 2018].

Salidroside, a phenylethanoid glycoside, consists of a tyrosol aglycone linked to a glucose moiety (p-hydroxyphenylethyl-O-β-D-glucopyranoside) [Tolonen et al., 2003]. It is widely distributed across *Rhodiola* species and is commonly used as a chemical marker for species identification [Dimpfel et al., 2018]. Salidroside has been reported to exhibit adaptogenic, antioxidant, anti-inflammatory, and anti-stress properties, along with regulatory effects on cellular energy metabolism [Li et al., 2021]. Despite these reported activities, the extent to which salidroside directly contributes to antioxidant capacity, particularly relative to other *Rhodiola* constituents, remains incompletely understood.

Phenylpropanoids such as rosavin, rosarin, and rosin are characteristic constituents of *Rhodiola rosea* and *Rhodiola sachalinensis*, with rosavin representing the major phenylpropanoid constituent [Dimpfel et al., 2018]. Rosavin (cinnamyl-(6-O-α-L-arabinopyranosyl)-O-β-D-glucopyranoside) is a cinnamyl-alcohol derivative containing a disaccharide composed of glucose and arabinose [Tolonen et al., 2003]. It has been associated with antioxidant, lipid-lowering, analgesic, antiradiation, antitumor, and immunomodulatory effects [Wang et al., 2023]. Structurally, rosavin lacks a free aromatic hydroxyl group commonly present in classical *para*-hydroxylated phenolic antioxidants; however, its phenylpropanoid backbone contains an extended conjugated system that permits electron delocalization, potentially enabling modest radical stabilization [Chen et al., 2015]. The presence of the disaccharide moiety may further influence its redox behavior as glycosidic substitution generally diminishes the capacity of hydroxyl groups to directly participate in redox chemistry [Wang et al., 2024]. Both rosavin and salidroside have been extensively studied for their adaptogenic, neuroprotective, anti-inflammatory, and antioxidant activities [Chiang et al., 2015]. Much of this bioactivity is attributed to their underlying chemical structures, namely, the phenylpropanoid or tyrosol backbone and associated glycosidic linkages, which influence polarity, metabolism, and redox behavior [Olennikov et al., 2020]. Several studies demonstrate that *Rhodiola* constituents can modulate oxidative stress pathways [Chiang et al., 2015; Kosakowska et al., 2018]; however, the degree to which these effects arise from direct radical scavenging versus indirect biological signaling remains unclear. The present study investigated antioxidant properties of the *Rhodiola rosea* extract and its chemical constituents to provide mechanistic insight into their biological activity.

## Materials and Methods

### Chemicals

*Rhodiola* Root Standardized Fluid Extract (Alcohol-Free) was from Nature’s Answer (Hauppauge, NY, USA). Rosavin (Cat# A424145) and salidroside (Cat# A130275) were purchased from Ambeed (Buffalo Grove, IL, USA). Arabinose (Cat# T13752), rosin (Cat# T4S0398), cinnamyl alcohol (Cat# T5S1550), and rosavin (Cat# T3886) were purchased from TargetMol (Wellesley Hills, MA, USA). Catalase from bovine liver (Cat# C-9322), 5,5′-dithiobis(2-nitrobenzoic acid) (DNTB; Cat# D8130), and L-cysteine (Cat# C-7755) were from Sigma-Aldrich (St. Louis, MO, USA).

### Assay kits to monitor reactive oxygen species (ROS)

EpiQuik In-Situ and Ex-Situ Hydrogen Peroxide (H_2_O_2_) Assay Kit (Cat# P-6005) was purchased from EpigenTek (Farmingdale, NY, USA). The fluorescence signals produced through the oxidation of a fluorogen, 10-Acetyl-3,7-dihydroxy-phenoxazine (Amplex Red) by H_2_O_2_ were measured using a POLARstar OPTIMA Fluorescence Microplate Reader (BMG Labtech, Cary, NC, USA). Amplite Colorimetric Hydrogen Peroxide Assay Kit (Cat# 11500) was purchased from AAT Bioquest (Pleasanton, CA, USA). Oxidation product of Amplite IR, exhibiting an intense blue color, was measured at an absorbance of 650 nm using an EMax Plus Microplate Reader (Molecular Devices, LLC., San Jose, CA, USA). Insta-Test Peroxide Test Strips Low Range (Cat# 2984LR) were purchased from LaMotte (Chestertown, MD, USA). Test strips were inserted into the reaction mixtures for 1 min at room temperature, photographed, and the intensities of the H_2_O_2_ signals were analyzed using ImageJ (National Institutes of Health, Bethesda, MD, USA). H_2_O_2_ levels were also assessed by detecting the fluorescence of the Europium-Tetracycline-H_2_O_2_ complex [Rakicioglu et al., 1999; Durkop & Wolfbeis, 2005] at an excitation wavelength of 390 nm and emission wavelength of 620 nm using a POLARstar OPTIMA Fluorescence Microplate Reader. H_2_O_2_ levels in rat serum and cell culture medium samples were measured using the QuantiChrom Peroxide Assay Kit (Cat# DIOX-250) purchased from BioAssay Systems (Hayward, CA, USA) on an EMax Plus Microplate Reader. Superoxide anion radicals produced by xanthine and xanthine oxidase were measured using the Amplite Colorimetric Superoxide Dismutase (SOD) Assay Kit (Cat# 11305) purchased from AAT BioQuest. A colored product generated from ReadiView SOD560 upon reaction with superoxide was monitored at the absorbance of 562 nm using an EMax Plus Microplate Reader.

### Molecular Dynamics Simulation

The minimum distance between H_2_O_2_ and rosavin, arabinose, or rosin, as well as the number of hydrogen bonds (H-bonds) formed between rosavin, arabinose, or rosin and water (H_2_O) or H_2_O_2_ were estimated using the GROMACS molecular dynamics simulation package (www.gromacs.org). Prior to the simulation, molecular docking was carried out using AutoDock Vina. The simulation was carried out at 300K temperature and 1 atm pressure.

### 5,5′-Dithio-bis-(2-nitrobenzoic acid) (DTNB) assay

Oxidation of the sulfhydryl group of L-cysteine was monitored using DTNB (Ellman’s Reagent) as previously described [Ellman, 1959; Suzuki et al., 1990]. DTNB reacts with free sulfhydryl groups to produce 2-nitro-5-thiobenzoate (TNB), which is a yellow-colored product quantified at an absorbance of 412 nm using an EMax Plus Microplate Reader.

### Cell culture experiments

Human pulmonary artery smooth muscles were purchased from ScienCell Research Laboratories (Carlsbad, CA, USA) and cultured in accordance with the manufacturer’s instructions in 5% CO2 at 37°C. Passages 3-6 were used. The number of viable cells was assessed using the Cell Counting Kit-8 (CCK8; Dojindo Molecular Technologies, Rockville, MD, USA).

### Animal experiments

Male Fischer CDF rats (Charles River Laboratories, Wilmington, MA, USA) were injected intraperitoneally with 0.4 mg/kg body weight of recombinant SARS-CoV-2 spike protein S1 subunit (Cat# 40591-V08H; Sino Biological, Paoli, PA, USA). Some rats were also injected with rosavin (32 mg/kg body weight twice a week). After 4 weeks, serum samples were collected for analysis. The Georgetown University Animal Care and Use Committee approved all animal experiments. Our investigation conformed to the National Institutes of Health Guide for the Care and Use of Laboratory Animals.

### Statistical Analysis

Means and standard errors of mean (SEM) were computed. Statistical analyses were performed using one-way analysis of variance (ANOVA). P < 0.05 was defined to be statistically significant.

## Results

### Rhodiola rosea inhibits H_2_O_2_

To characterize the antioxidant properties of *Rhodiola rosea* and its components, the effects of *Rhodiola rosea* extract on H_2_O_2_ levels were tested using a 10-acetyl-3,7-dihydroxy-phenoxazine fluorogen-based EpiQuik H_2_O_2_ Assay. As shown in Fig. 1A, the addition of H_2_O_2_ to the assay resulted in increased fluorescence signal. In the presence of *Rhodiola rosea* extract, H_2_O_2_ signals were attenuated, suggesting that the extract reduces H_2_O_2_ levels. This effect of *Rhodiola rosea* was dose-dependent (Fig. 1B). Importantly, the *Rhodiola rosea* extract alone did not affect the fluorescence-based H_2_O_2_ assay (Fig. 1A); however, the intrinsic brown color of the extract did not allow for assessment by colorimetric assays.

**Figure 1:**
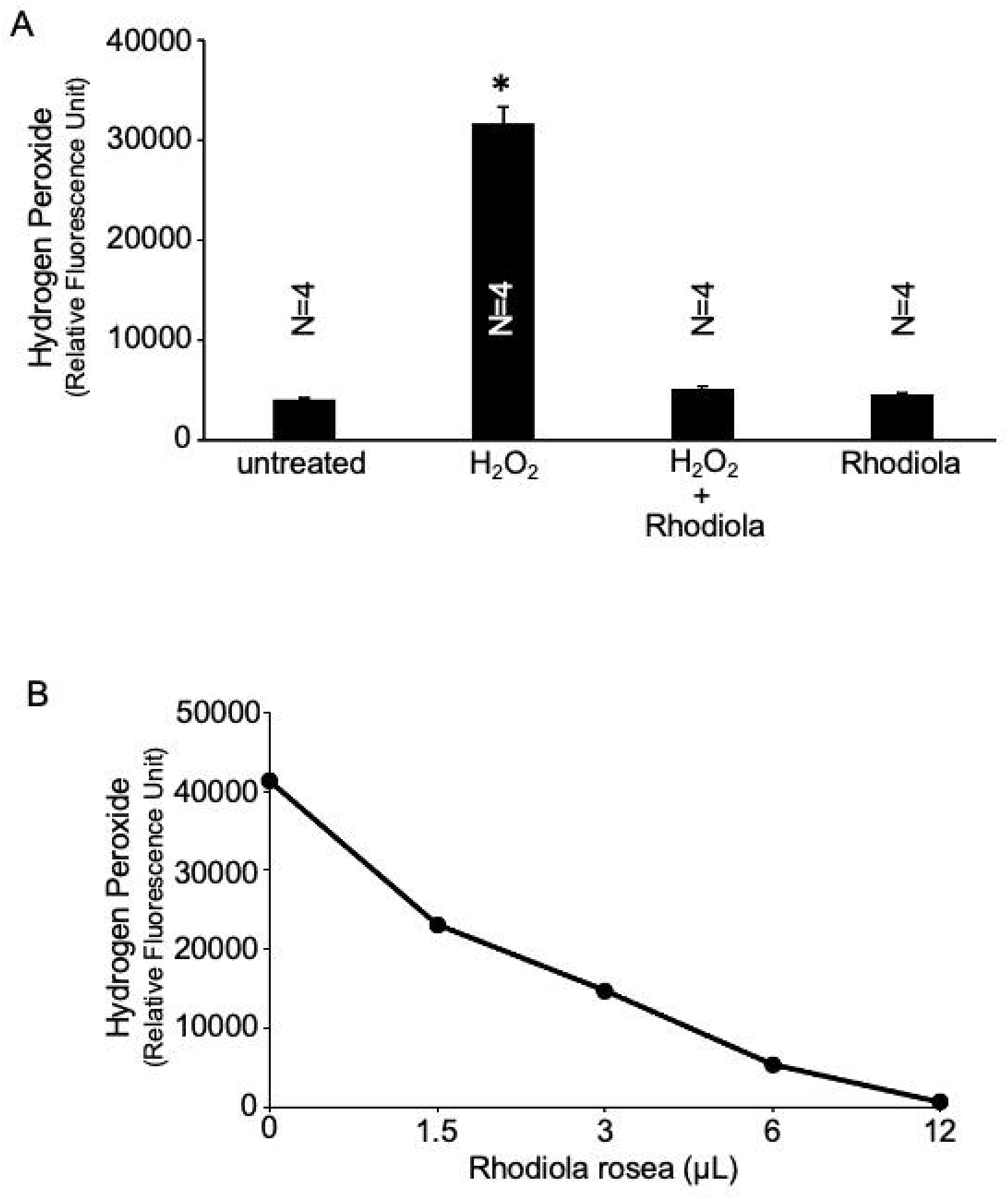
Effects of *Rhodiola rosea* on H_**2**_**O**_2_. (A) EpiQuik In-Situ and Ex-Situ Hydrogen Peroxide (H_2_O_2_) Assay (EpigenTek) was used to determine the effects of the *Rhodiola rosea* extract on H_2_O_2_. The assay reactions contained a 4 μM final concentration of H_2_O_2_ along with 6 μL *Rhodiola rosea* extract. The bar graph represents mean ± SEM. * indicates that the value is significantly different from other groups at p < 0.05 as determined by One-way ANOVA. (B) Dose-response of the effects of various doses of Rhodiola rosea on a 2.5 μM final concentration of H_2_O_2_.

### Rosavin inhibits H_2_O_2_

Given that the major constituents of *Rhodiola rosea* are rosavin and salidroside, the effects of these compounds on H_2_O_2_ levels were evaluated. Initial experiments using the Amplite Colorimetric H_2_O_2_ Assay showed that rosavin alone did not interfere with assay readout. Fig. 2A demonstrates that the H_2_O_2_ signal was significantly attenuated by a positive control, catalase, as well as rosavin; however, salidroside did not have an effect. The inhibitory effects of rosavin on H_2_O_2_ were found to be dose-dependent as determined by both the absorbance Amplite Assay (Fig. 2B) and the fluorescence-based EpiQuik Assay (Fig. 2C). In this study, rosavin purchased from Ambeed was used for all the experiments with the exception of Fig. 2D, which shows that rosavin from another source (TargetMol) also exhibited the effects on H_2_O_2_. Consistent results on the inhibitory effects of rosavin were also obtained using other detection methods, including the Lamotte Insta-Test Hydrogen Peroxide Test Strips (Fig. 3A) and Europium-Tetracycline assay (Fig. 3B).

**Figure 2:**
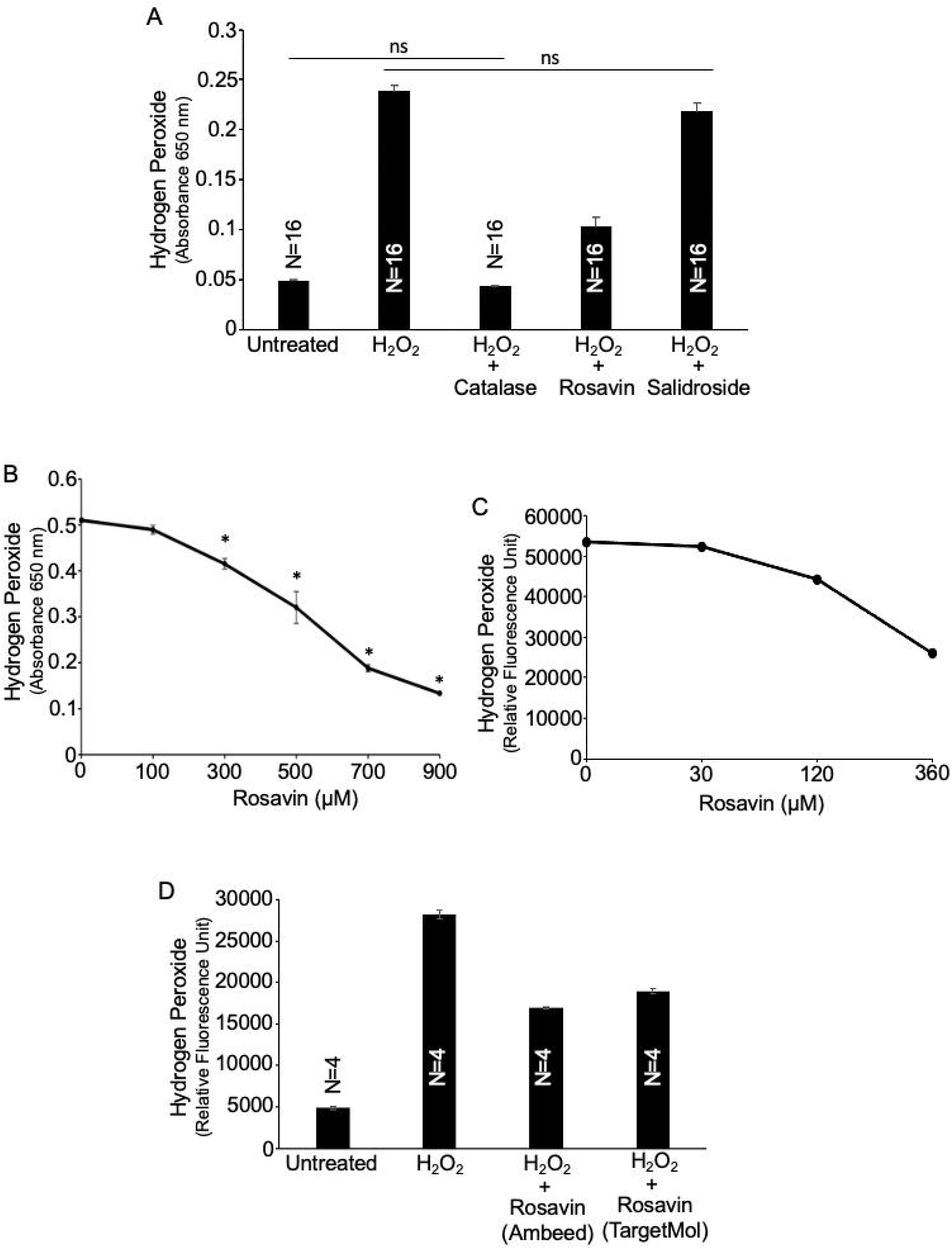
Rosavin, but not salidroside, inhibits H_**2**_**O**_2_. (A) Amplite Colorimetric Hydrogen Peroxide Assay (AAT Bioquest) was used to determine the effects of rosavin and salidroside on H_2_O_2_. The assay reactions contained a 10 μM final concentration of H_2_O_2_, catalase (1,000 U/mL), which is a known H_2_O_2_ scavenger, rosavin (400 μM), and/or salidroside (400 μM). The bar graph represents mean ± SEM. ns indicates not significantly different from each other. All the values without ns denotation are significantly different at p < 0.05 as determined by One-way ANOVA. (B) Rosavin produced a concentration-dependent decrease in H_2_O_2_ signal, as determined by the Amplite Colorimetric Hydrogen Peroxide Assay. The assay reactions contained a 10 μM final concentration of H_2_O_2_ and various concentrations of rosavin. The line graph represents mean ± SEM (n = 4). * denotes groups that are significantly different from 0 μM rosavin control at p < 0.05 as determined by One-way ANOVA. (C) Rosavin produced a concentration-dependent decrease in H_2_O_2_ signal, as determined by the EpiQuik In-Situ and Ex-Situ Hydrogen Peroxide Assay. The assay reactions contained a 4 μM final concentration of H_2_O_2_ along with various concentrations of rosavin. (D) The EpiQuik In-Situ and Ex-Situ Hydrogen Peroxide Assay shows that the effects of rosavin (obtained from Ambeed), as shown above, were also reproduced by rosavin from another source (TargetMol).

**Figure 3:**
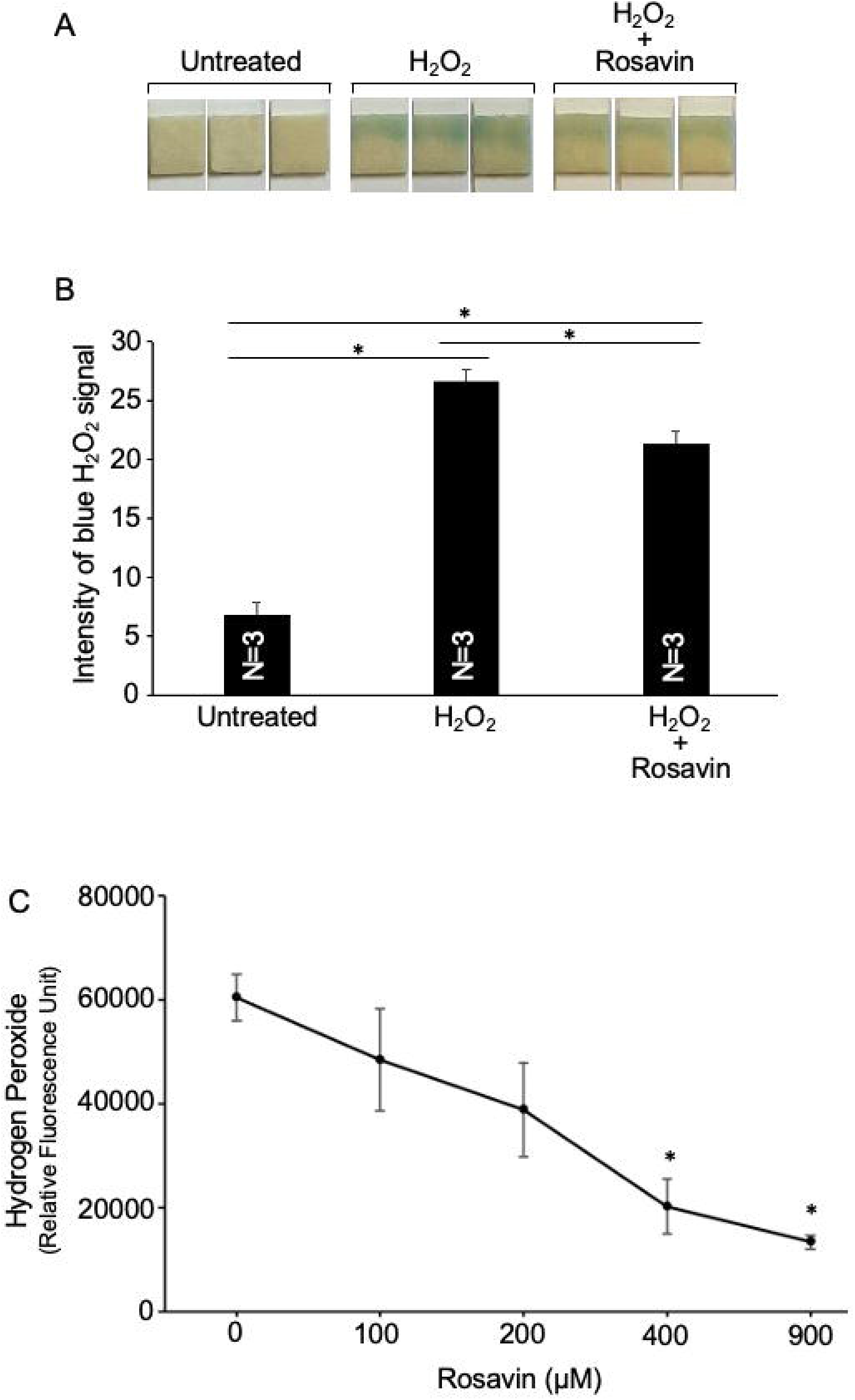
Rosavin inhibits H_**2**_**O**_**2**_ as assessed by Peroxide Strips and Europium-Tetracycline Assay. (A) Reaction mixtures contained a 10 μM final concentration of H_2_O_2_ and 400 μM rosavin. Insta-Test Peroxide Test Strips, Low Range (LaMotte, Chestertown, MD, USA) were inserted into the reaction mixture for 1 min at room temperature. Representative results of Peroxide Strips. (B) The bar graph represents mean ± SEM values obtained from image analysis using Image J. * denotes groups that are significantly different from each other at p < 0.05 as determined by One-way ANOVA. (C) Europium-Tetracycline assay was used to determine the effects of rosavin on H_2_O_2_ levels. The line graph represents mean ± SEM (n = 5). * indicates that the value is significantly different from 0 μM rosavin control at p < 0.05 as determined by One-way ANOVA.

By contrast, rosavin and salidroside did not influence the superoxide anion radical levels, as monitored using the Amplite Colorimetric Superoxide Assay. While the positive control, superoxide dismutase (SOD), effectively decreased the superoxide signal, neither rosavin nor salidroside exerted any effects (Fig. 4).

**Figure 4:**
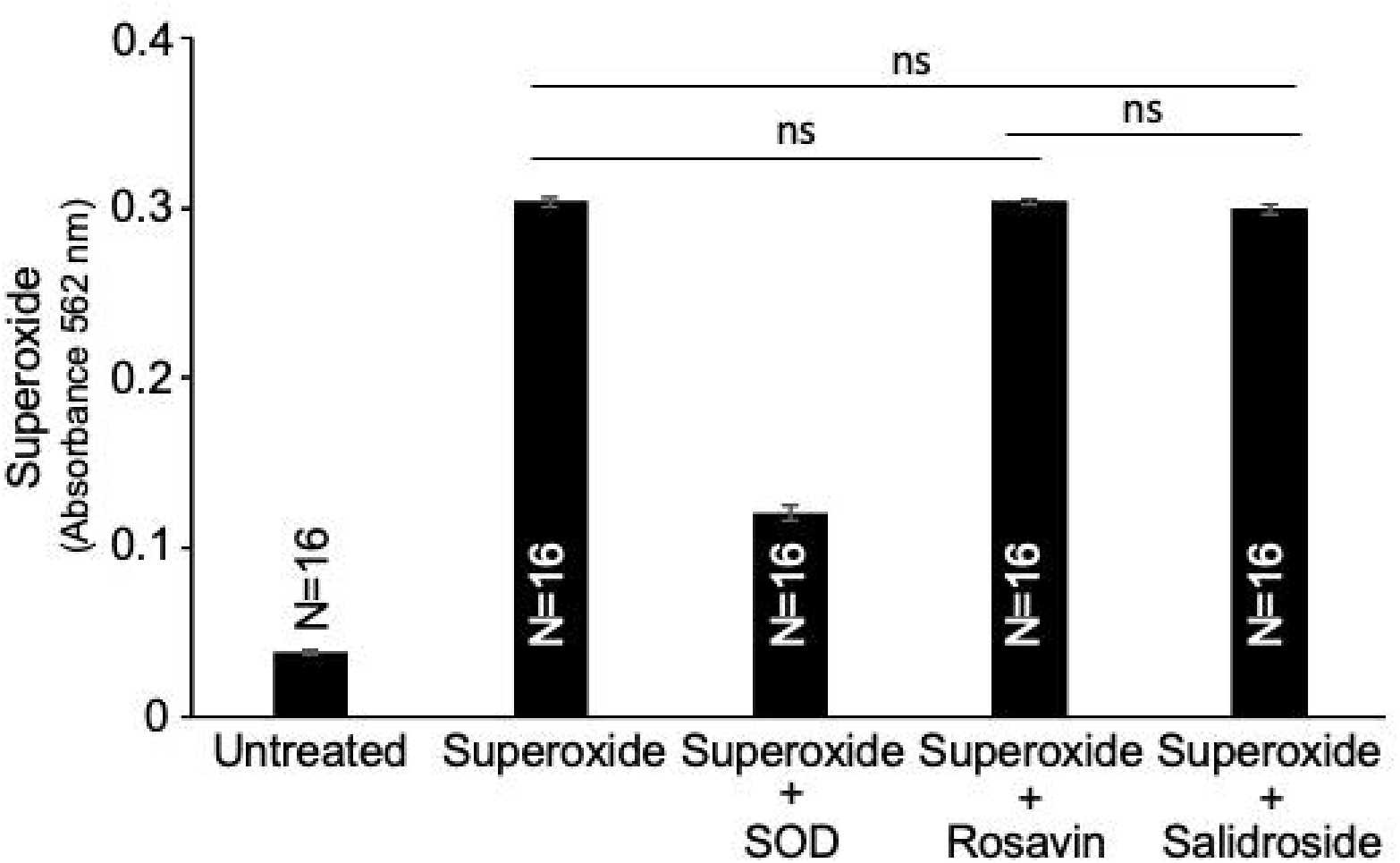
Neither rosavin nor salidroside exerts effects on superoxide anion radical. Amplite Colorimetric Superoxide Dismutase (SOD) Assay (AAT Bioquest) was used to determine the effects of rosavin and salidroside on superoxide produced by xanthine plus xanthine oxidase. Superoxide was detected using ReadiView SOD560. The assay reactions contained xanthine plus xanthine oxidase, SOD (5 U/mL), which is a known superoxide scavenger, rosavin (400 μM), and/or salidroside (400 μM). The bar graph represents mean ± SEM. ns indicates not significantly different from each other. All the values without ns denotation are significantly different at p < 0.05 as determined by One-way ANOVA.

To investigate the mechanism underlying rosavin-mediated inhibition of H_2_O_2_, we performed structure-function studies to determine which structural features are necessary for activity. Rosavin is composed of arabinose (a monosaccharide) linked to rosin (a monoglucoside derivative of cinnamyl alcohol). Using the Amplite Colorimetric H_2_O_2_ Assay, we found that arabinose (Fig. 5A), rosin (Fig. 5B), and cinnamyl alcohol (Fig. 5B) did not exhibit H_2_O_2_ inhibitory activity comparable to rosavin. Results from the EpiQuik Fluorescence H_2_O_2_ Assay confirmed these observations (Fig. 5C). We further tested higher concentrations of arabinose and rosin (750 μM) to ensure that these chemical components do not have the ability to reduce H_2_O_2_ (Fig. 5D). Furthermore, the addition of arabinose and rosin together did not influence H_2_O_2_ levels (Fig. 5C).

**Figure 5:**
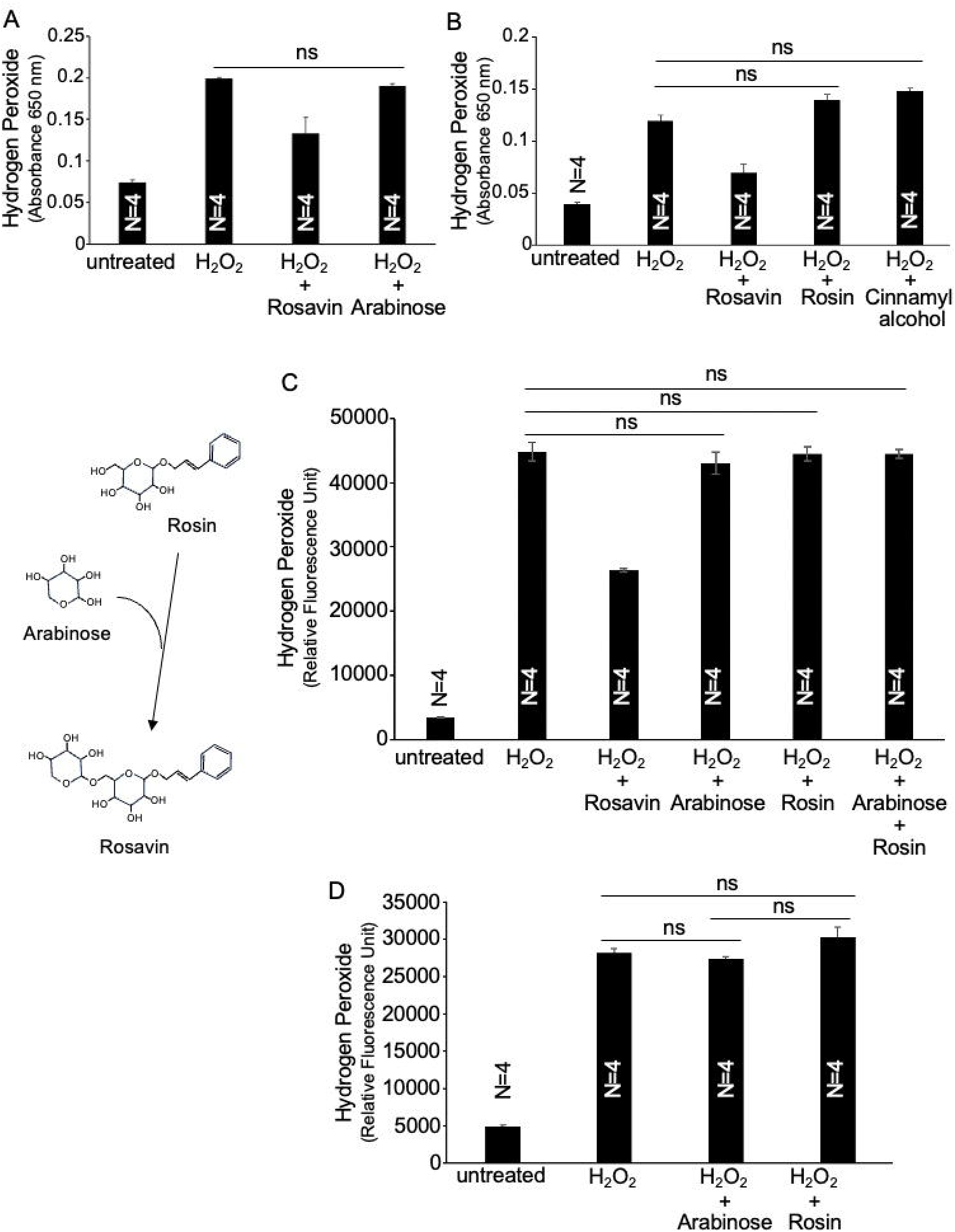
Structure-function relationship studies. (A and B) Ampliate Colorimetric Hydrogen Peroxide Assay (AAT Bioquest) and (C and D) EpiQuik In-Situ and Ex-Situ Hydrogen Peroxide Assay (EpigenTek) were used to determine the effects of rosavin, arabinose, and rosin on H_2_O_2_. The assay reactions contained a 3 μM final concentration of H_2_O_2_ and rosavin, arabinose, rosin, and/or cinnamyl alcohol at 150 μM. (D) Effects of arabinose or rosin (750 μM) were also tested to ensure that the results showing the inability of these molecules to influence H_2_O_2_ are not because the amounts tested were too small. The bar graphs represent mean ± SEM. ns indicates not significantly different from each other. All the values without ns denotation are significantly different at p < 0.05 as determined by One-way ANOVA.

Molecular dynamics simulations demonstrated that rosavin exhibited a shortest minimum distance to H_2_O_2_ compared to arabinose or rosin (Table 1). The results also showed that water formed more H-bonds with rosavin compared to rosin or arabinose, whereas H-bonding between H_2_O_2_ and any of the compounds (rosavin, arabinose, or rosin) was negligible (Table 1).

**Table 1:** Molecular Dynamics Simulation.

| Solute | Minimum Distance<br>to H<sub>2</sub>O<sub>2</sub> (nm) | Number of H-Bonds<br>between solute and H<sub>2</sub>O<sub>2</sub> | Number of H-Bonds<br>between solute and H<sub>2</sub>O |
| --- | --- | --- | --- |
| Rosavin | 1.83 | 0.22 ± 0.008 | 11.12 ± 0.074 |
| Arabinose | 2.11 | 0 | 7.69 ± 0.060 |
| Rosin | 2.75 | 0 | 7.34 ± 0.066 |

Values represent means ± SEM.

Together, these results indicate that the intact glycoside structure of rosavin defines its ability to inhibit H_2_O_2_.

### Functional roles of rosavin inhibition of H_2_O_2_

To assess the functional consequences of rosavin-mediated inhibition of H_2_O_2_, we first examined whether rosavin could attenuate H_2_O_2_-induced oxidation of thiol groups. 5,5′-dithio-bis-(2-nitrobenzoic acid), DTNB or Ellman’s Reagent, reacts with free sulfhydryl groups to produce 2-nitro-5-thiobenzoate (TNB), a yellow-colored product that can be detected at an absorbance of 412 nm (Ellman, 1959). As shown in Fig. 6, L-cysteine containing a reduced sulfhydryl group reacted with DTNB to produce this signal, which was diminished upon oxidation by H_2_O_2_. Addition of rosavin in this reaction effectively prevented H_2_O_2_-mediated oxidation of cysteine thiol groups, confirming rosavin’s capacity to preserve thiol redox status through reduction of H_2_O_2_.

**Figure 6:**
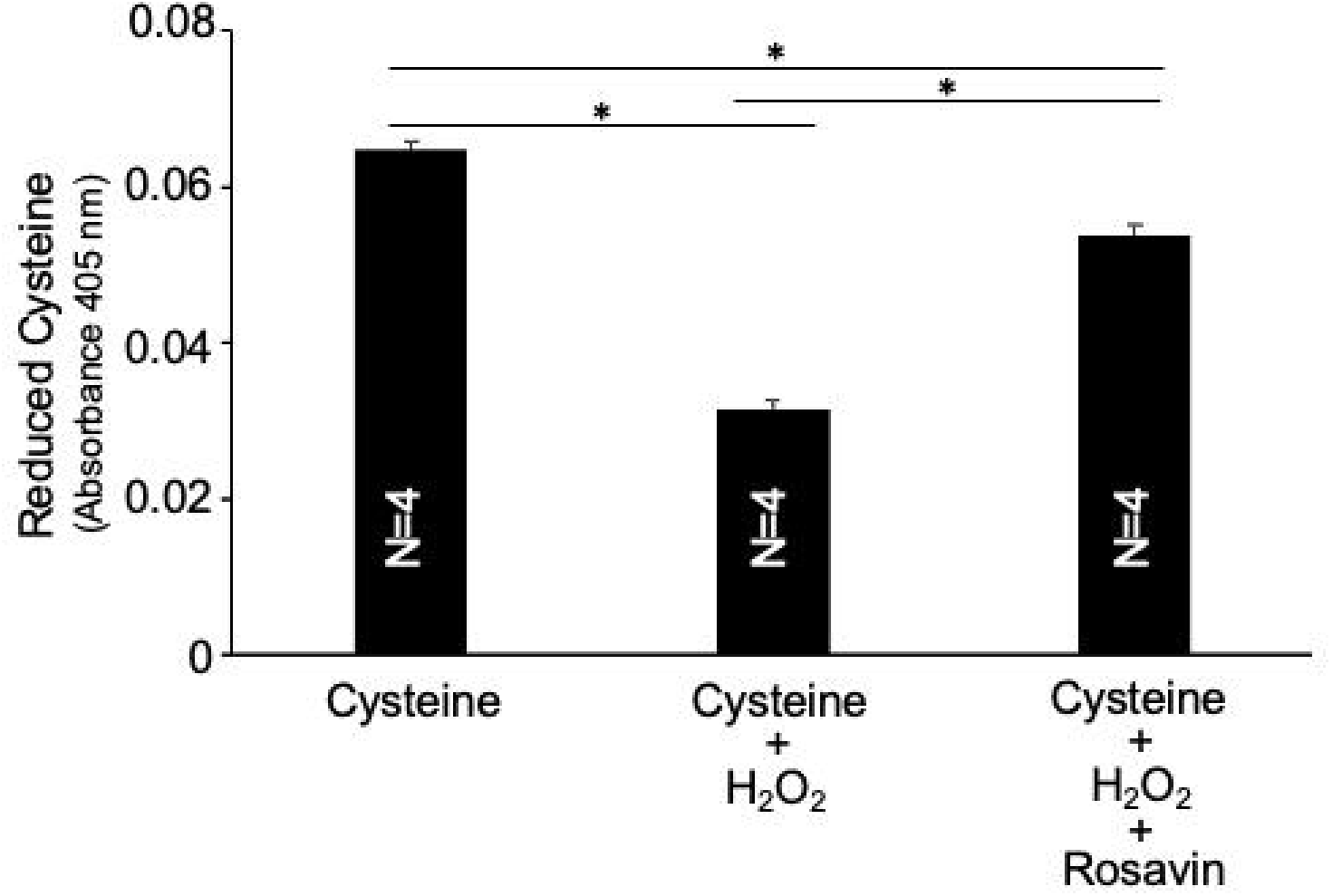
Rosavin inhibits H_**2**_**O**_**2**_-mediated oxidation of thiols. DTNB was used to monitor reduced thiol content. Reaction mixtures contained 0.5 mM DTNB, 1 mM L-cysteine, 25 μM H_2_O_2_, and 1 mM rosavin. The bar graph represents mean ± SEM. * denotes groups that are significantly different from each other at p < 0.05 as determined by One-way ANOVA.

We next examined whether rosavin could mitigate H_2_O_2_-induced effects in cultured cells. The treatment of human vascular smooth muscle cells with H_2_O_2_ decreased cell viability, as monitored by the CCK-8 assay, which measures cellular metabolic activity via the reduction of the water-soluble tetrazolium salt (WST-8) by dehydrogenases. Treatment with rosavin attenuated the H_2_O_2_-induced reduction in cell viability, supporting a cytoprotective role of rosavin during oxidative stress (Fig. 7A). In this cell culture model, we also observed that rosavin decreased the levels of H_2_O_2_ released into the culture media (Fig. 7B).

**Figure 7:**
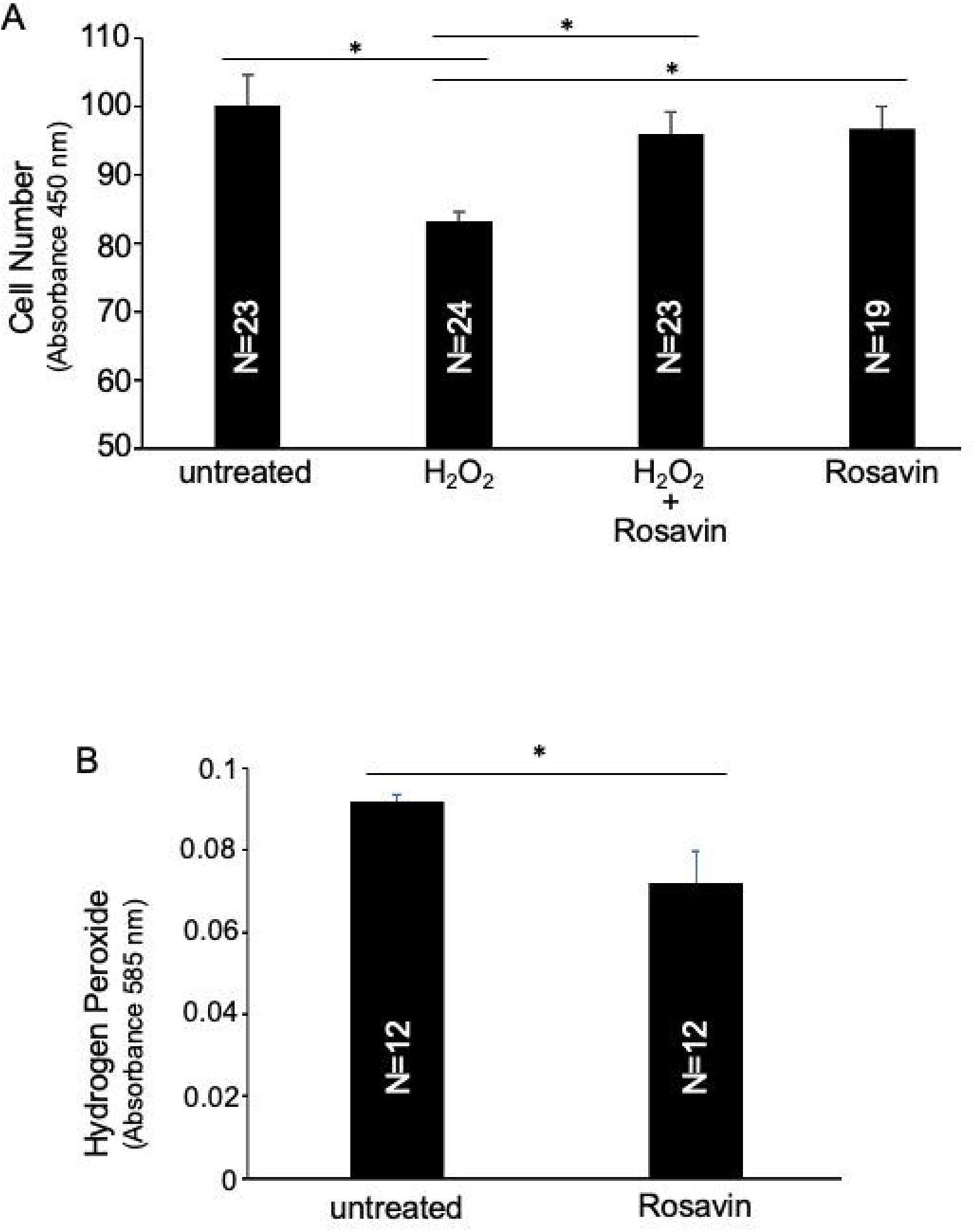
Effects of rosavin on H_**2**_**O**_**2**_ in cultured cells. Cultured human pulmonary artery smooth muscle cells were treated with 1 μM H_2_O_2_ and 40 μM rosavin for 2 days. (A) The cell number of viable cells was determined by CCK8 assay, demonstrating rosavin inhibited H_2_O_2_-effects on cells. (B) H_2_O_2_ levels were measured in culture media, demonstrating that rosavin decreased H_2_O_2_ levels using QuantiChrom Peroxide Assay. Bar graphs represent mean ± SEM. * denotes groups that are significantly different from each other at p < 0.05 as determined by (A) One-way ANOVA and (B) Student’s t-test.

Finally, we investigated the effects of rosavin *in vivo* using Fischer 344 rats injected with the SARS-CoV-2 spike protein S1 subunit. Circulating H_2_O_2_ levels were increased 4 weeks after the intraperitoneal injection of the S1 subunit, demonstrating enhanced systemic oxidative stress. Treatment of these rats with rosavin by intraperitoneal injection twice weekly for 4 weeks significantly decreased the levels of H_2_O_2_ (Fig. 8).

**Figure 8:**
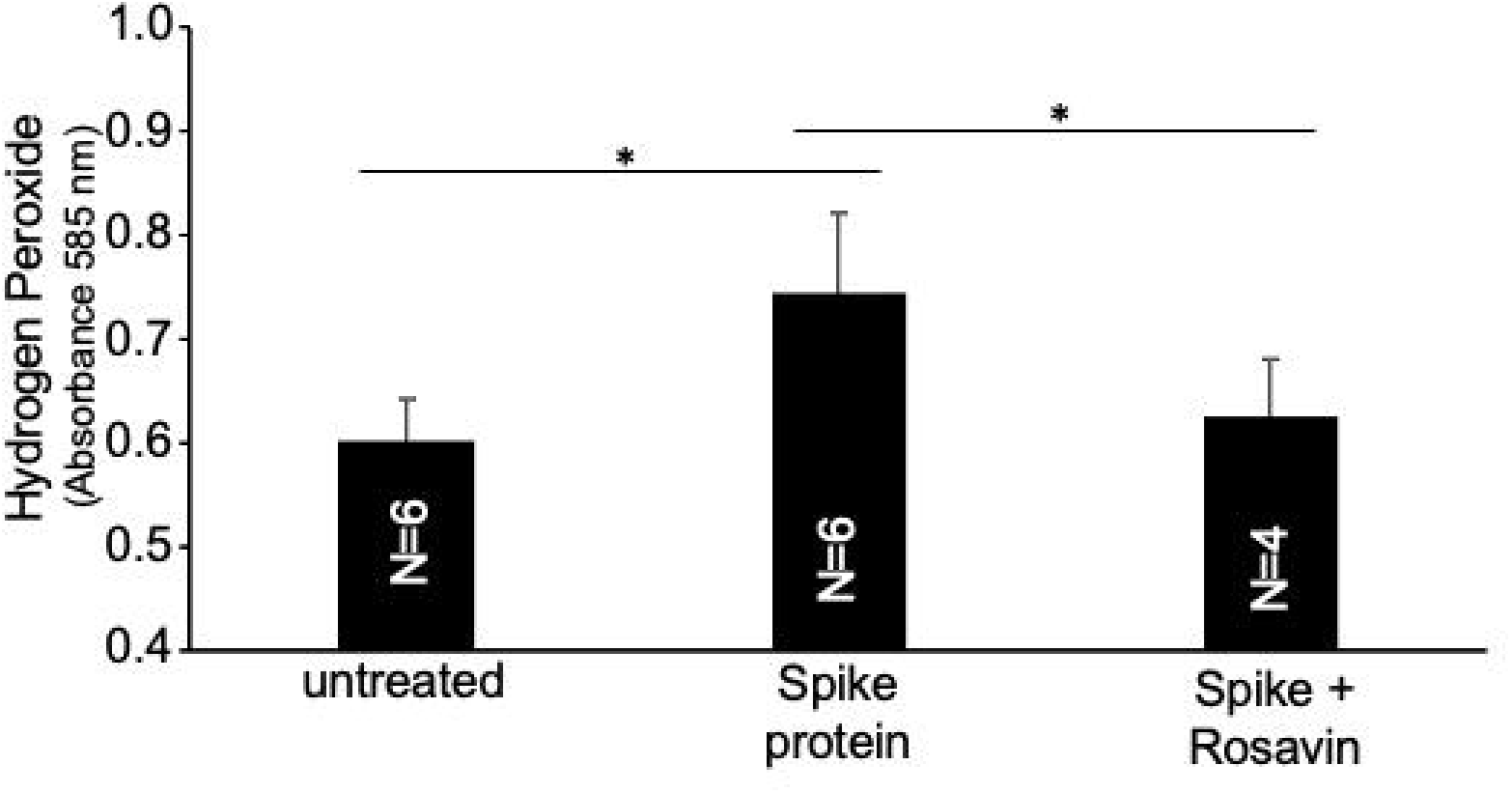
Rosavin administration reduces circulating H_**2**_**O**_2_ in rats. Fischer (CDF) rats were intraperitoneally injected with the SARS-CoV-2 spike protein S1 subunit. After 4 weeks, serum samples were obtained. Some rats were also subjected to intraperitoneal injection of rosavin twice a week for 4 weeks. Serum H_2_O_2_ levels were measured using QuantiChrom Peroxide Assay. The bar graph represents mean ± SEM. * denotes groups that are significantly different from each other at p < 0.05 as determined by One-way ANOVA.

## Discussion

The major finding of the present study is that *Rhodiola rosea* extract and its major constituent, rosavin, reduce H_2_O_2_ levels in *in vitro* biochemical assays, cultured human cells, and in *in vivo* rat models. These findings provide mechanistic insights into the antioxidant and cytoprotective properties attributed to *Rhodiola rosea* extract and identify rosavin as a structurally dependent bioactive component contributing to these effects.

In this study, rosavin, but not salidroside, consistently attenuated H_2_O_2_ signals across several H_2_O_2_ detection assays. This pattern was further supported by the cysteine/DTNB assay, where rosavin attenuated H_2_O_2_-mediated oxidation of cysteine thiol groups. In contrast, neither rosavin nor salidroside reduced superoxide in the xanthine/xanthine oxidase assay, indicating a lack of detectable electron-transfer reactivity toward superoxide. Together, these findings suggest that rosavin displayed a measurable chemical reactivity with H_2_O_2_, while neither rosavin nor salidroside displayed detectable reactivity with superoxide under the conditions tested.

Rosavin contains a cinnamyl alcohol derived phenylpropanoid backbone, which consists of an aromatic ring conjugated to an unsaturated three-carbon side chain [Grech-Baran et al., 2014]. Such extended conjugation allows for partial electron delocalization, which stabilizes radical or cationic intermediates that may form during redox interactions (Szeląg et al., 2014). While rosavin does not possess the *para*-hydroxyl group usually associated with phenolic antioxidants, its conjugated phenylpropanoid scaffold itself can still support limited hydrogen atom transfer or single electron transfer under certain chemical conditions (Wang et al., 2024). These structural features provide a plausible explanation for rosavin’s ability to chemically reduce H_2_O_2_. While glycosidic substitution may reduce direct interaction of the aromatic system with oxidants, the underlying cinnamyl alcohol motif appears sufficient to support modest radical stabilization [Tominaga et al., 2005]. Nevertheless, our experimental results indicate that the intact rosavin structure is required to exert inhibitory activity toward H_2_O_2_ (Fig. 5).

The ability of the intact rosavin structure to reduce H_2_O_2_ may be related to its shorter minimum distance to H_2_O_2_, compared to arabinose or rosin (Table 1). Additionally, rosavin was not predicted to form H-bonds with H_2_O_2_, suggesting that direct H-bond interactions between rosavin and H_2_O_2_ are unlikely to contribute to the observed effect. The simulation results also demonstrated that rosavin formed more H-bonds with surrounding water molecules than arabinose or rosin, raising the possibility that stronger interactions with the aqueous environment may enhance the efficacy of contact with H_2_O_2_ and facilitate the antioxidant reaction.

The degree of conjugation differs markedly between rosavin and salidroside. Salidroside’s tyrosol backbone consists of an aromatic ring attached to a saturated ethyl chain, lacking the extended π-system present in rosavin [Liang et al., 2024]. As a result, salidroside has a reduced capacity to delocalize charge or stabilize radical intermediates, making electron or hydrogen donation less favorable [Liang et al., 2024]. This structural limitation of salidroside likely explains its lack of detectable interaction with H_2_O_2_ in purely chemical reactions [Kosakowska et al., 2018].

*In vivo* experiments further demonstrated that administration of the spike protein S1 subunit of Severe Acute Respiratory Syndrome Coronavirus 2 (SARS-CoV-2), which caused the COVID-19 pandemic [Huang et al., 2020] and continues to affect millions of people who suffer from long COVID [de Melo et al., 2025], increased H_2_O_2_ levels in rats. These levels were decreased by rosavin treatment. These observations suggest that rosavin may mitigate oxidative stress in a physiologically relevant context. While the present study focuses specifically on the relationship between H_2_O_2_ and rosavin, ongoing work in our laboratory is rigorously investigating the broader pathologic outcomes elicited by the spike protein exposure and the potential therapeutic role of rosavin as a strategy to treat COVID-19-associated pathologies.

In summary, this study provides mechanistic insights into how *Rhodiola rosea* and its constituents may influence redox biology and highlights a structure-dependent role of rosavin in modulating H_2_O_2_. These findings contribute to a more detailed understanding of the functional properties of *Rhodiola*-derived compounds.

